# Interstage flow matrices: population statistic derived from matrix population models

**DOI:** 10.1101/2023.06.14.545022

**Authors:** Hiroyuki Yokomizo, Keiichi Fukaya, John G. Lambrinos, Yuka Kawai, Takenori Takada

## Abstract

1. Many population statistics describe the characteristics of populations within and among species. These are useful for describing population dynamics, understanding how environmental factors alter demographic patterns, testing hypotheses related to the evolution of life history characteristics, and informing the effective management of populations.
2. In this study, we propose a population statistic: the interstage flow. The interstage flow is defined as the product of the element in the *i*th row, the *j*th column of the population projection matrix, and the *j*th element of the normalized stable stage distribution.
3. The sum of the interstage flow matrix elements is equal to the population growth rate, which is the dominant eigenvalue of the population projection matrix. The interstage flow matrix elements allow decomposition of population growth rate into component contributions made by transitions between developmental stages.
4. We demonstrate the utility of interstage flow matrices using matrix population models from the COMPADRE plant matrix database. We compared interstage flows among four life history/functional groups (semelparous herbs, iteroparous herbs, shrubs, and trees) and described how population growth rate reflected individual transitions related to stasis, fecundity, and growth. We found that the individual flows are different among functional groups.
5. *Synthesis:* The proposed population statistic, the interstage flow matrix, describes the contribution of individual developmental stage transitions to the population growth rate. The flow of individuals between developmental stages differs in distinctive ways among different life histories and functional groups. The interstage flow matrix is a valuable statistic for describing these differences.

## INTRODUCTION

Detailed demographic data have been compiled by plant ecologists for populations that span a range of taxonomic, life history, and environmental conditions. Researchers have developed a number of statistics to describe these data, and the development of convenient population statistics has been an important aspect of theoretical population ecology. Matrix population models (MPMs) can be used to derive many useful population statistics, such as population growth rate, generation time, age at maturation, sensitivity, and elasticity. These statistics are helpful for describing population dynamics, understanding how environmental factors alter demographic patterns, and testing hypotheses related to the evolution of life history characteristics (Silvertown et al., 1993; Silvertown et al., 1996; Salguero-Gómez et al., 2016; Salguero-Gómez, 2017).

The growing availability of plant demographic data has allowed more robust hypothesis tests and spurred innovative new directions in comparative plant demography. For example, Silvertown et al. (1996) calculated MPM elasticities of about 90 plant species and showed that the relative influence of different developmental stage transitions (e.g., recruitment from seed, stasis, transitions to larger size classes) on population growth varied systematically across plant functional groups. These differences likely reflect morphological, physiological, and environmental constraints that influence the evolution of plant life history. In 2015, the COMPADRE database began compiling published plant MPMs in a standardized and accessible format. Salguero-Gómez et al. (2016) used COMPADRE to quantitatively describe the life history strategies of 418 plant species explicitly in terms of their demographic characteristics. They found that species assorted along two relatively independent strategies: a slowly growing, long-lived strategy and a reproduction-focused strategy.

New statistics that describe demographic properties from different perspectives could be useful for answering certain questions. In this study, we introduce a statistic called the interstage flow, which is expressed in matrix form and describes the flow of individuals between developmental stages or ages. The approach provides information that is not provided by standard population statistics based on the population projection matrix such as population growth rate, sensitivity, and elasticity. The elements of a population projection matrix reflect vital rates—i.e., survival, growth, and fecundity—of individuals within demographic classes. They do not explicitly describe the demographic contribution of the individuals transitioning between classes. The flow of individuals between classes is a distinct aspect of demography. For example, a particular developmental stage might have high seed production, but if the proportional abundance of individuals in that stage is low, its contribution to population growth rate may be minimal. In contrast to the population projection matrix, the matrix elements of interstage flow explicitly decompose population growth into the contributions made by individuals transitioning between different developmental stages.

The idea of interstage flow was brought by Kawano et al. (1987), but they neither formally interpreted its ecological meaning nor comprehensively analyzed its properties. Also, no work has been done to test how interstage flow matrices vary across taxa or functional groups. In this study we formally define the interstage flow matrix, describe some of its useful properties, compare it to related statistics such as elasticity, and interpret its ecological meaning. We examine how variation in interstage flow is related to plant functional form across 286 plant species using data from the COMPADRE database.

## MATERIALS AND METHODS

### Definition of interstage flow

The dynamics of populations over a discrete time interval can be described as follows:

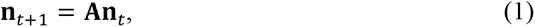

where **A** is the population projection matrix, and **n***_t_* indicates a vector of population sizes of each stage at time *t*. We propose a metric (statistic): the interstage flow matrix **F_IS_**. It is defined as follows:

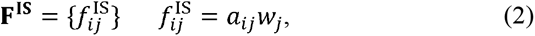

where ***a****_ij_* is a matrix element of **A**. *w_j_* is the *j*th element of the stable stage distribution (**w**) of matrix **A**. The stable stage distribution is normalized such that the sum of all elements is equal to 1, that is,

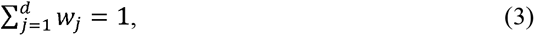

where *d* denotes the matrix dimension of **A**.

We present how to obtain the interstage flow matrix in Figure 1 using a population projection matrix of a Japanese perennial herb, *Trillium apetalon* (Ohara et al., 2001), as an example. The population projection matrix **A** is

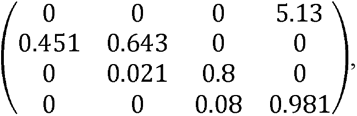

and the normalized stable stage distribution can be obtained from the matrix as:

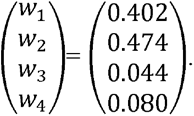

Therefore, the (2, 1) element of the interstage flow matrix is 0.451 × 0.402 (= 0.181) and the (1, 4) element of the interstage flow matrix is 5.13 × 0.08 (=0.410). These are not the probabilities of transitions between stages, but rather the normalized number of individuals transitioning between stages. Then, the interstage flow matrix is

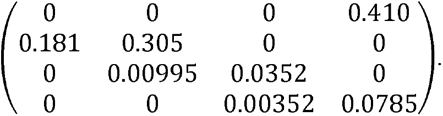

**Fig. 1.**
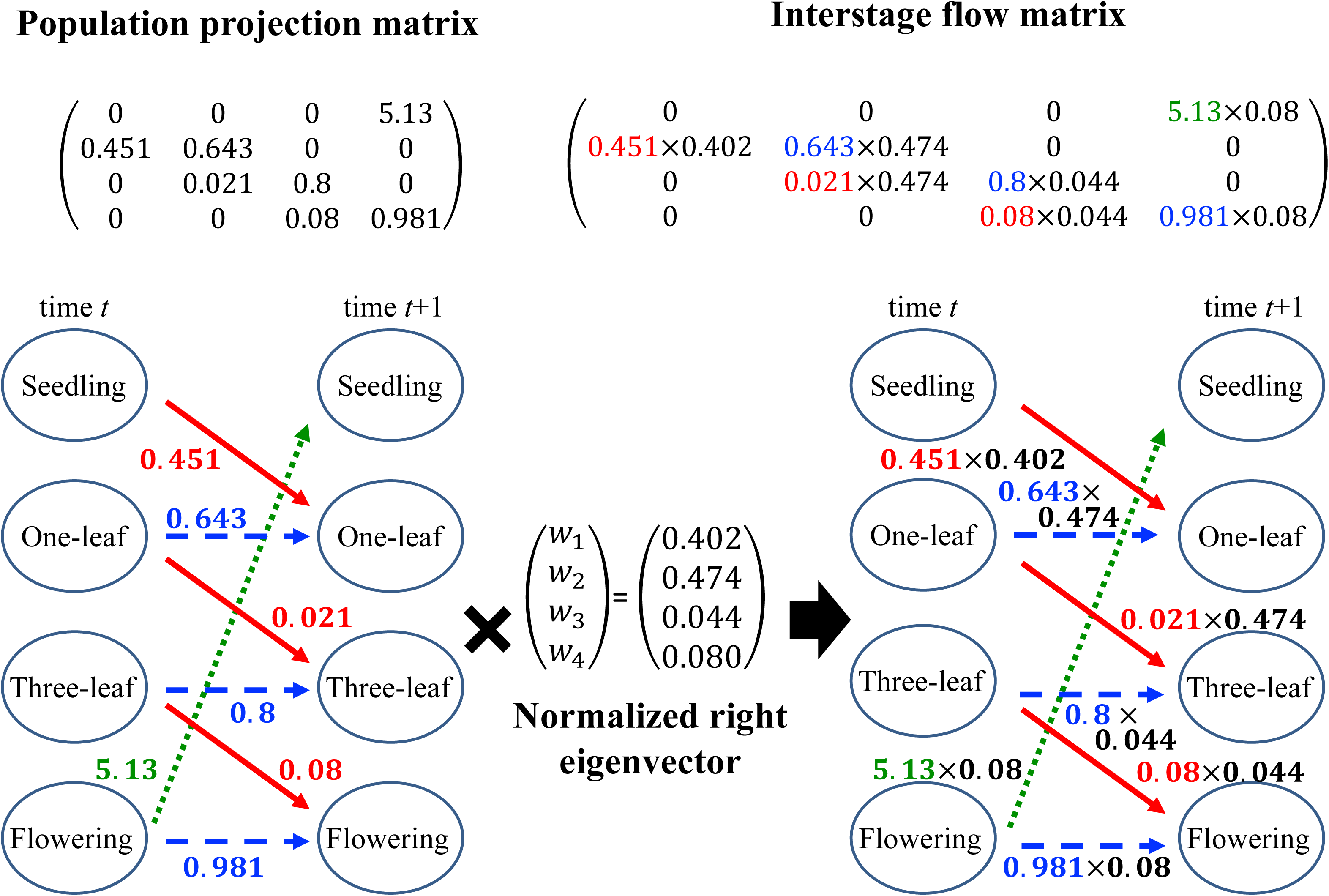
The transition probabilities between stages and interstage flow of the Japanese perennial herb, *Trillium apetalon*. Four stages (seedling, one-leaf, three-leaf, and flowering) were set to construct the population projection matrix of the species. The population projection matrix and the interstage flow matrix are shown on the top of the figure. The matrix is from Ohara et al. (2001) and was constructed using long-term census data of the Japanese perennial herb, *Trillium apetalon*. The flow chart between stages is shown on the bottom of the figure. The numbers attached to each arrow in the flow chart are the transition probabilities between stages. The normalized stable stage distribution (***w***) in the center was obtained from the population projection matrix. Using the stable stage distribution and the population projection matrix elements, the interstage flows between stages were calculated as shown in the right part of the figure (see Equation 2). Interstage flows are shown by each arrow on the right and in the interstage flow matrix.

The unique property of *f_ij_*^IS^ is that the relationship between the population growth rate λ and the elements of the flow matrix is as follows:

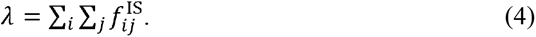

Equation 4 implies that the elements of the interstage flow matrix quantify the extent to which each transition between developmental stages contributes to the population growth rate.

If matrix **A** is primitive, it is confirmed by the strong ergodic theorem that the stable stage distribution (the vector of population size at time *t*) is:

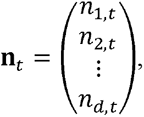

which is proportional to the normalized stable stage distribution (**w**):

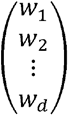

i.e.,

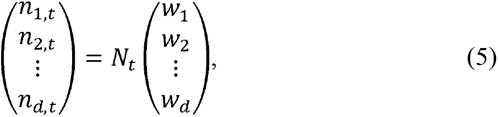

where 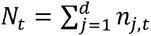.

During a single timestep, *N_t_w_j_* individuals, the number of individuals at stage *j,* processes of survival and reproduction. During the survival process, *N_t_w_j_* individuals flow to several stages, including stage *j*. These flows reflect the demographic flow to stage *i* with probability *a_ij_*, and the number decreases because *a_ij_* is less than 1. The amount of interstage flow from stage *j* to stage *i* (the number of survivors) is *N_t_a_ij_w_j_*. In the reproduction process, *N_t_w_j_* individuals reproduce their offspring with per-capita fecundity *a_ij_*. The amount of interstage flow in the reproduction process is similar to that in the survival process, *N_t_a_ij_w_j_*. The sum of all interstage flows is 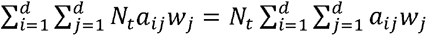, which must be equal to *N_t+_*_1_ because the sum of the surviving individuals and new offspring at the next time step is *N_t_*_+1_.

This can be proven by the following mathematical calculations: The number of *n_j,t_* increases or decreases as a single time step proceeds. This was calculated using Equation 1 as:

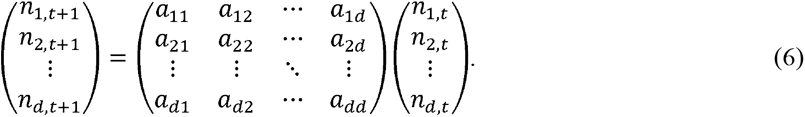

Equation 6 can be written based on Equation 5 as follows:

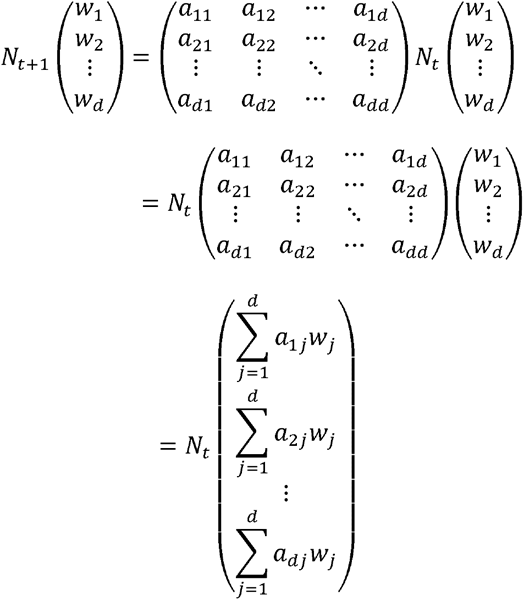

The sum of the elements of the left-hand side of the equation is *N_t_*_+1_, and the sum of the elements of the right-hand side is 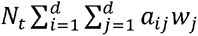. It is supposed that the population dynamics described by Equation 1 grow at a population growth rate and with a structure proportional to the stable stage distribution after sufficient time has elapsed (p. 86 in Caswell [2001]). Hence, *N_t_*_+1_ = *λN_t_*. Therefore, 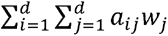 = *λ*. This means that all interstage flow matrix elements sum to the population growth rate. We provide another proof of Equation 4 in Appendix A.

### Elasticity analysis

Elasticity is frequently used to quantify relative change in the population growth rate λ per relative change in matrix elements (Caswell et al., 1984; de Kroon et al., 1986; Pfister, 1998; Kaneko & Takada, 2014; Yokomizo et al., 2017; Takada & Kawai, 2020). Interstage flow and elasticity are related as follows (see Appendix B for details):

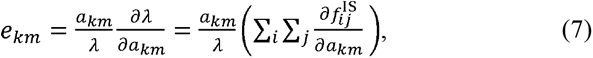

where *e_km_* and *f_ij_*^IS^ are the elasticity and flow matrix elements, respectively. Equation 7 indicates that the sensitivity *δλ*/*δa_km_* is the sum of changes in interstage flows when the matrix element changes.

### COMPADRE plant matrix database

We described how interstage flows (Equation 2) are related to life history and population characteristics for a set of plant populations derived from the COMPADRE database version 6.21.6. (Salguero-Gómez et al., 2015; www.compadredb.org). The COMPADRE database contains matrix population models of 758 vascular plant species consisting of four matrices: population projection matrix **A** and submatrices **U**, **F**, and **C (A** = **U** + **F** + **C)** (see Fig. 2a). The submatrix **U** contains the transition and survival rates of individuals in each age or stage. Submatrix **F** contains the number of seeds produced per individual, and submatrix **C** contains the clonal reproduction rates.

**Fig. 2.**
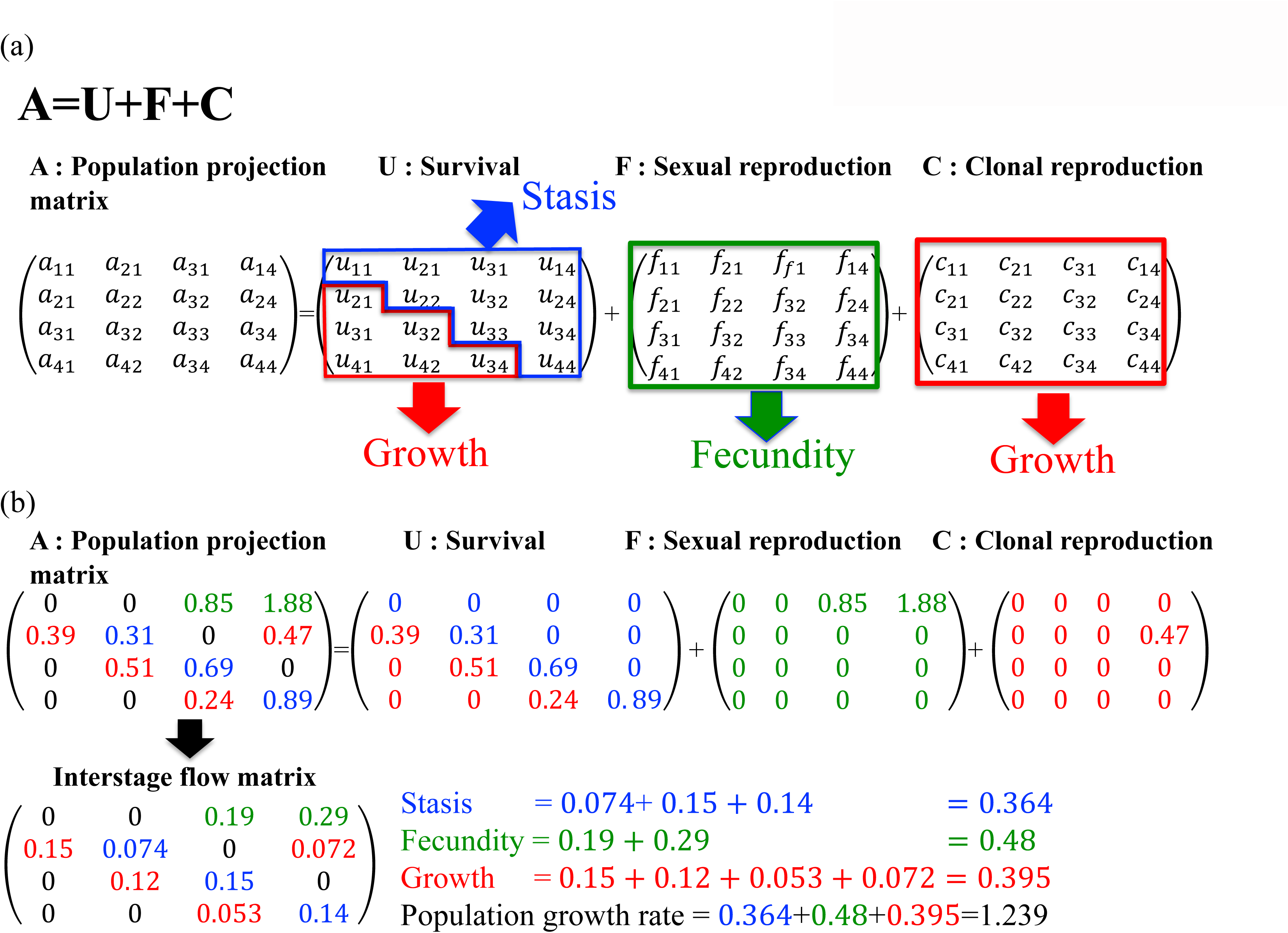
Classification of matrix elements into stasis, fecundity, and growth. (a) Population projection matrix **A** is partitioned into three submatrices: **U**) contains only transitions and survival of existing individuals; **F**) sexual reproduction; **C**) clonal reproduction. The diagonal or upper diagonal elements of **U** were classified as “stasis”, and lower diagonal elements of **U** were classified as “growth”. The elements of **F** and **C** were classified as “fecundity” and “growth”, respectively. (b) A hypothetical example of matrix elements partitioned into stasis, fecundity, or growth. The classification for each element is identical between population projection matrix and flow matrix. Flow matrix elements are summed for each class (i.e., stasis, fecundity, and growth). The sum of the flow matrix elements is the population growth rate.

### Species functional classification

We categorized the selected matrices into the following growth form categories: herbaceous perennials, shrubs, and trees based on the “OrganismType” classification in the metadata of the COMPADRE database. We also categorized herbaceous perennials as either semelparous or iteroparous. Semelparous herbs die immediately after reproduction and typically have high fecundity to compensate for the loss of future reproductive opportunities (Charnov & Schaffer, 1973; Pianka, 1976, 1978). The trade-off between fecundity and adult survival in semelparous herbs affects elasticity (Takada & Kawai, 2020). Semelparous herbs were identified from the elements of population projection matrices by using the method described by Takada et al. (2018).

### Matrix selection

We used matrices satisfying the following eight criteria:

1. No maximum stage-specific survival in the submatrix U exceeded one.
2. At least one element in submatrices **F** or **C** was greater than zero.
3. Matrices were irreducible and primitive. This is because we considered that the population density in a stage does not become zero or oscillate between years if the stage distribution is stable. A matrix is irreducible or primitive if (**I**+**A)***^d^*^-1^ is positive and 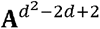 is positive, respectively, where *d* is the dimension of the population projection matrix of **A** (Caswell, 2001).
4. Dimensions were equal to or larger than four because small matrices cannot reflect stage-specific information about transitions.
5. Matrices were for unmanipulated populations. We removed matrices for experimentally manipulated populations.
6. Matrices satisfied **A** = **U** + **F** + **C**. For some populations, this equation does not hold true, for various reasons (e.g., errors in the database).
7. Higher stages in matrices are for larger sizes, active or matured stages. In some matrices, dormant stages are in larger rows than active ones with the same size categories. We did not use those matrices in this study, because we assumed transitions to larger rows are classified as growth.
8. Only for analyses involving elasticity, we used matrices in which all matrix elements can be classified into only one category out of stasis, fecundity, and growth (See the next section).

When multiple matrices were available for a species, we calculated the mean matrix and the average of each element. However, we could not calculate a mean matrix if there were multiple population projection matrices with different dimensions. In such cases, we selected the population projection matrix associated with the study with the longest research period.

We obtained interstage flow matrices for 14 species of semelparous herbs, 143 species of iteroparous herbs, 36 species of shrubs, 93 species of trees, and 286 species in total. In addition, we obtained elasticity matrices for 14 species of semelparous herbs, 135 species of iteroparous herbs, 36 species of shrubs, 91 species of trees, and 276 species in total.

### Classification of matrix elements into stasis, fecundity, and growth

We classified each interstage flow matrix element as reflecting the developmental stage transitions of fecundity, growth to larger size classes, or stasis within the same stage using an approach similar to the method used by Silvertown et al. (1996) (see Fig. 2a). The diagonal or upper diagonal elements of **U** (transitions to the same or lower stages) were classified as stasis and lower diagonal elements of **U** (transitions to per individual) were classified as fecundity. All elements of **F** (clonal reproduction) larger stages) were classified as growth. All elements of **C** (number of seeds produced were classified as growth.

An example of the classification process is shown in Fig. 2b. In this case, each element is classified into stasis, fecundity, or growth. The flow matrix is derived according to Equation 2 using the population projection matrix **A**. We summed the elements of the flow matrix for stasis, fecundity, and growth, respectively (see Fig. 2b). To allow comparison of interstage flows among plant species with different population growth rates, we divided the stasis, fecundity, and growth by its population growth rate—defined as normalized stasis, fecundity, and growth respectively. The sum of the normalized stasis, fecundity, and growth equals 1.0.

### Statistical analysis

We calculated interstage flow matrices for the selected plant species based on Equation 2. We used Dirichlet regression to describe how the patterns of interstage flows vary across plant functional groups. Dirichlet regression is used when a set of bounded variables has a constant sum, for example with proportions and probabilities (Adler et al., 2014). In our study, the sum of interstage flows related to stasis, fecundity and growth normalized by population growth rate was one, making Dirichlet regression a suitable choice. Matrix dimension (the number of developmental stage transitions described by the matrix) is not an innate characteristic of plant species, but rather is defined by researchers and reflects the design and constraints of individual studies. Matrix dimension has been shown to influence other statistics derived from matrices, such as population growth rate and elasticity (Ramula and Lehtila, 2005; Ramula et al., 2008). We therefore included matrix dimension as an explanatory variable in our analysis. Categorical variables for functional groups (FG), population growth rate per year (PGR), matrix dimensions (MD), interaction terms of FG and PGR, interaction terms of FG and MD, and interaction terms of PGR and MD were included in our model as explanatory variables. All analyses used R version 4.1.0 software (R Core Team, 2021), and we used the DirichReg() function in the DirichletReg package (Maier, 2021).

## RESULTS

Functional groups had distinct patterns of flow allocation among stasis, fecundity, and growth (Table 1, Fig. 3, and Fig. 4). The ternary plots in Fig. 3 illustrate these differences. Semelparous herbs had interstage flows that tended to be dominated by growth and fecundity (Fig. 3a). The spatial median, which shows the central tendency of the distribution of interstage flows, of semelparous herbs is located in an area with large values of growth and fecundity, suggesting that the interstage flows of semelparous herbs tended to be dominated by growth and fecundity. At the other end of the spectrum, the interstage flows of trees tended to be dominated by stasis (Fig. 3d). Iteroparous herbs and shrubs tended to have intermediate patterns that spanned a range between those two extremes (semelparous herbs and trees) (Fig. 3).

**Fig. 3.**
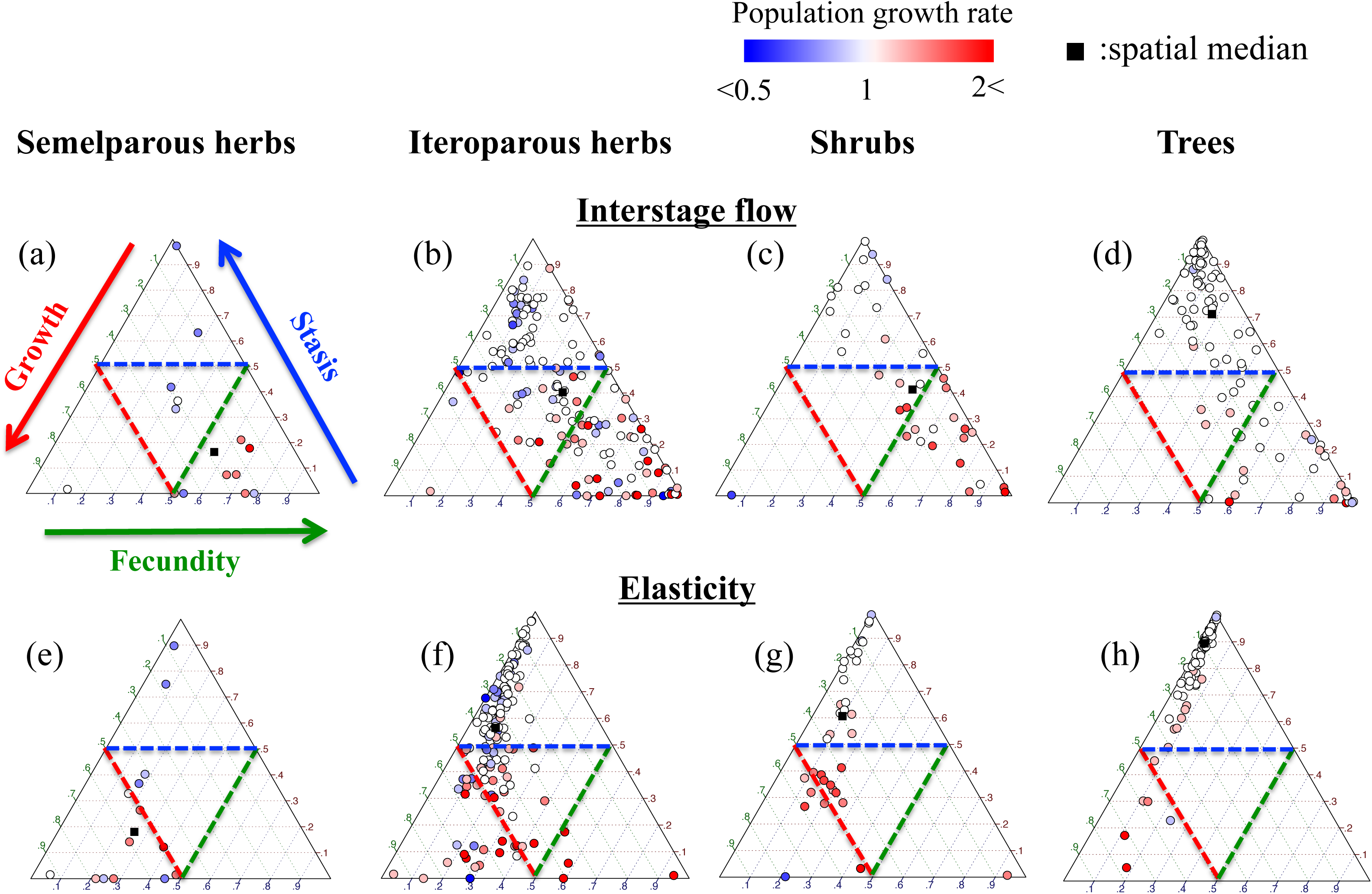
Ternary plot of interstage flow (a–d) and elasticity (e–h). Semelparous herbs, are plotted in (a) and (e). Iteroparous herbs are plotted in (b) and (f). Shrubs are plotted in (c) and (g). Trees are plotted in (d) and (h). Color shading indicates population growth rates. Dashed lines indicate a value of 0.5 for each axis. Filled squares indicate the spatial median (Vardi & Zhang, 2000).

**Fig. 4.**
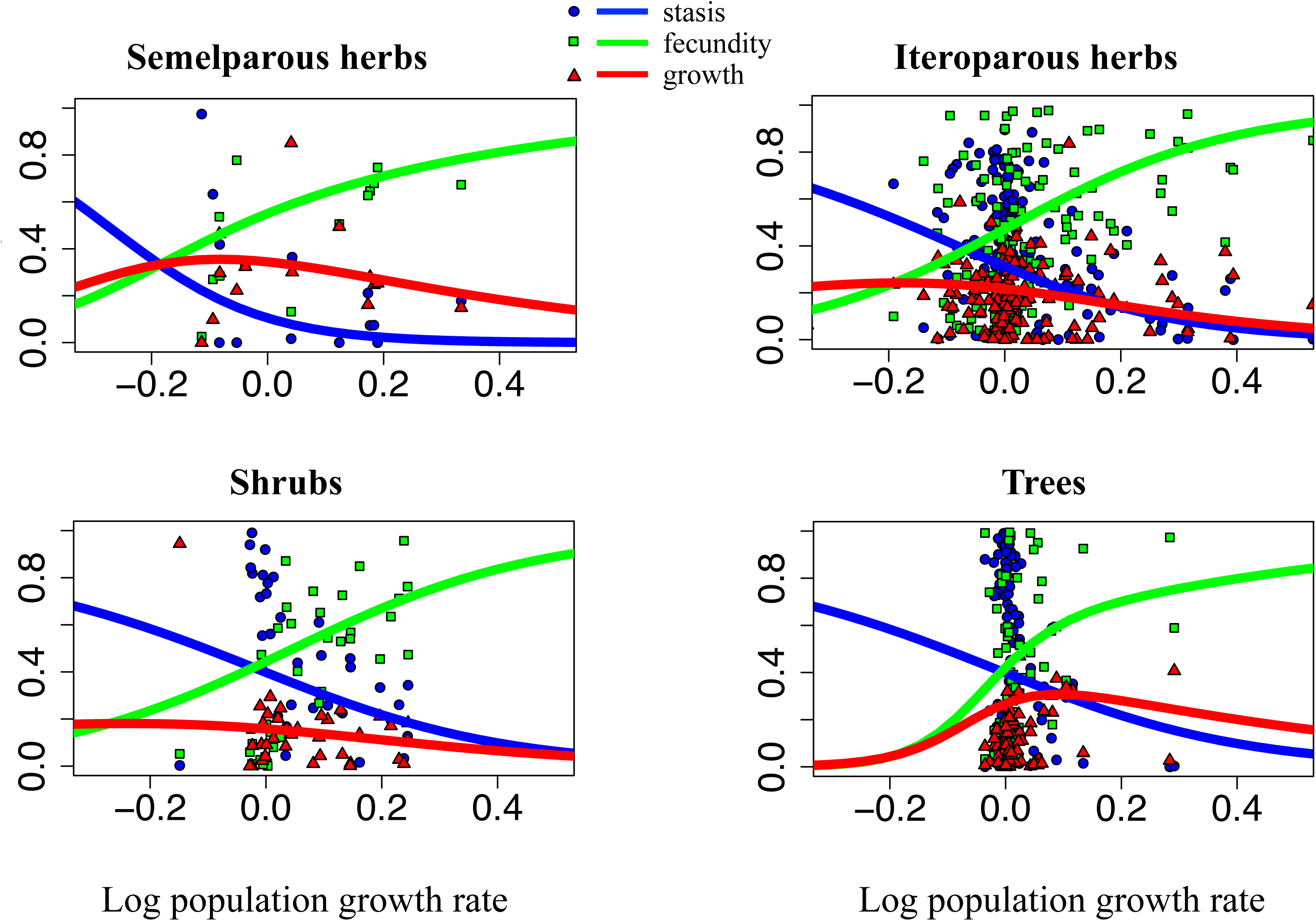
Dependence of interstage flow related to stasis, fecundity, and growth on the logarithmic population growth rate. (a) semelparous herbs, (b) iteroparous herbs, (c) shrubs, and (d) trees. The matrix dimensions used in the regression were the mean values of each functional group.

**Table 1.**
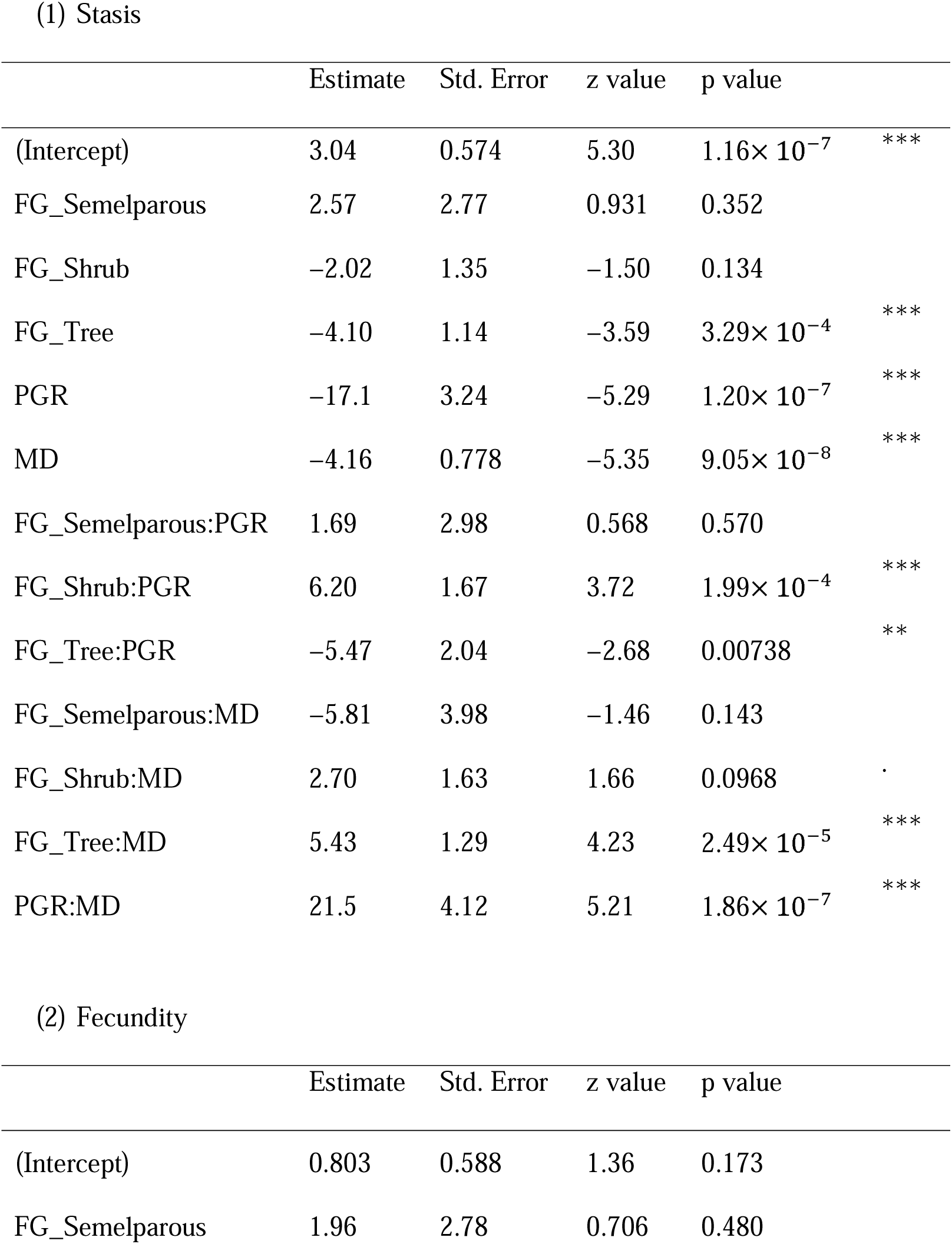
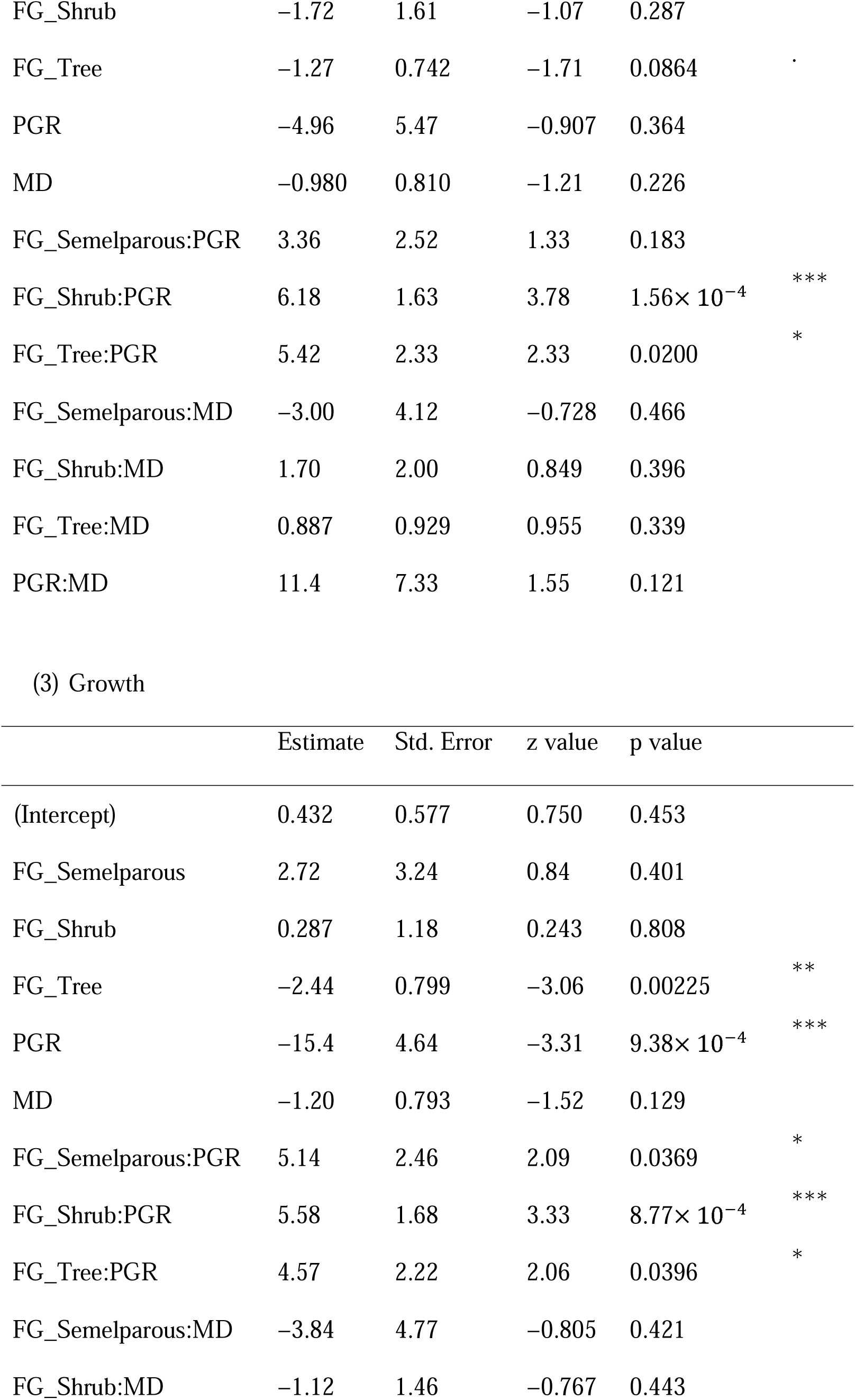
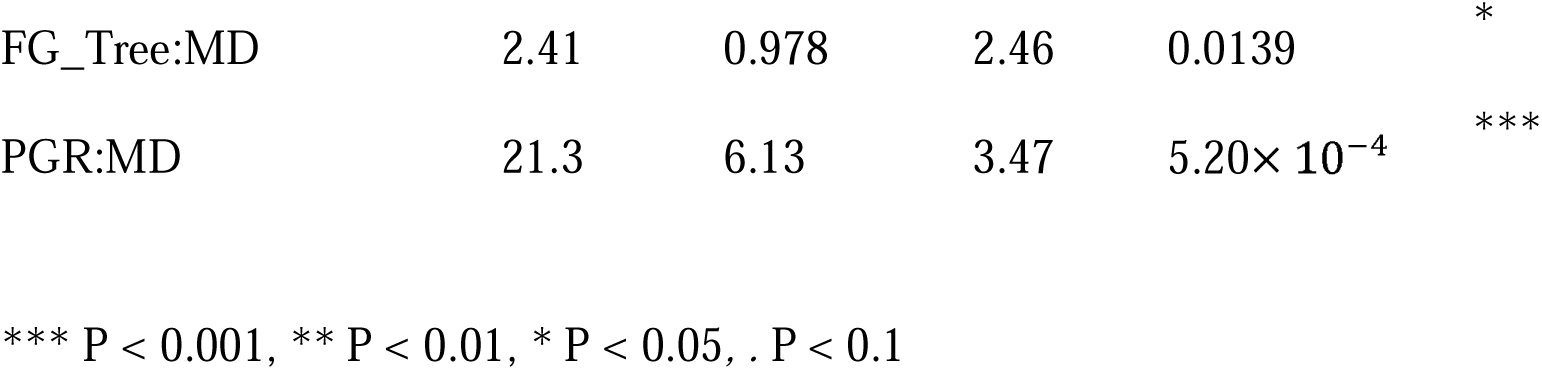
Summary of the results of Dirichlet regression for summed interstage flows for (1) stasis, (2) fecundity, and (3) growth. FG_Semelparous, FG_Shrub, and FG_Tree are categorical variables for semelparous herbs, shrubs, and trees, respectively. PGR is the population growth rate, and MD is the matrix dimension.

Population growth rate and matrix dimension also influenced patterns of interstage flow. Growing populations tended to have large interstage flows related to fecundity and relatively small flows related to both growth and stasis (Fig. 4). Overall, population growth rate had a statistically significant influence on interstage flows related to stasis and growth, but the influence differed among functional groups (see interaction terms between FG and PGR in Table 1). Interstage flows related to stasis tended to decrease with matrix dimension, but this influence also varied among functional groups and PGR (Table 1).

The pattern of elasticities among functional groups broadly mirrored those observed for interstage flow. Trees tended to have large elasticities for stasis, and semelparous herbs tended to have large elasticities for growth, whereas iteroparous herbs and shrubs had elasticities that spanned an intermediate range between growth and stasis. But in contrast to interstage flow, elasticities generally ascribed less of an influence on fecundity (Fig. 3). Also, in contrast to interstage flow, the largest elasticities in growing populations tended to be related to growth (Fig. 5). Overall, population growth rate, matrix dimension, and their interaction had a statistically significant influence on the elasticities related to stasis, fecundity, and growth (Table 2).

**Fig. 5.**
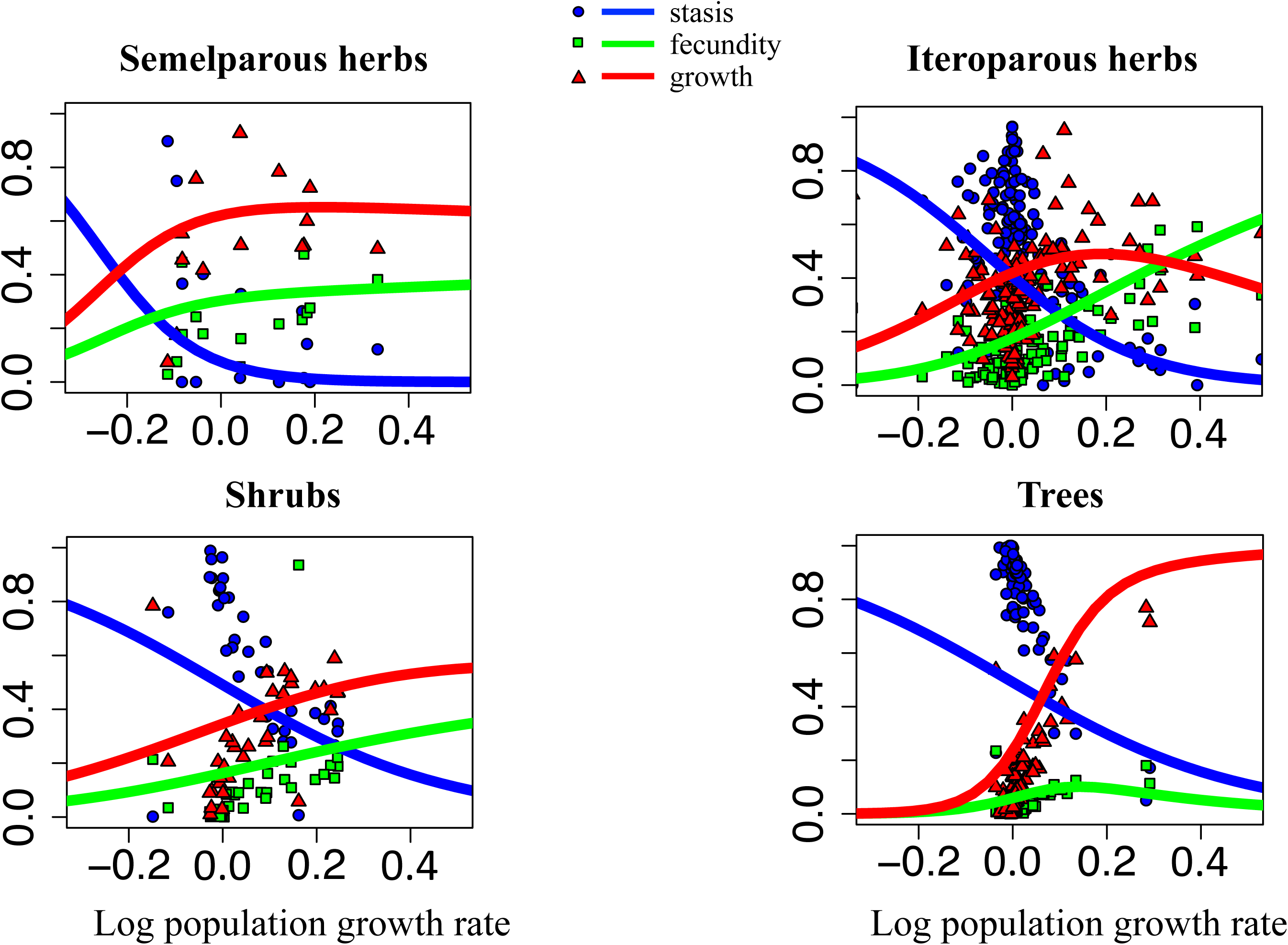
Dependence of elasticity related to stasis, fecundity, and growth on the logarithmic population growth rate. (a) semelparous herbs, (b) iteroparous herbs, (c) shrubs, and (d) trees. The matrix dimensions used in the regression were the mean values of each functional group.

**Table 2.** Summary of the results of Dirichlet regression for summed elasticities for (1) stasis, (2) fecundity, and (3) growth. FG_Semelparous, FG_Shrub, and FG_Tree are categorical variables for semelparous herbs, shrubs, and trees, respectively. PGR is the population growth rate, and MD is the matrix dimension.

|  | Estimate | Std. Error | z value | p value |  |
| --- | --- | --- | --- | --- | --- |
| (Intercept) | 5.11 | 0.485 | 10.5 | $<2.00 \times 10^{-16}$ | *** |
| FG_Semelparous | 1.32 | 2.56 | 0.514 | 0.607 |  |
| FG_Shrub | -2.7 | 1.64 | -1.65 | 0.0997 | . |
| FG_Tree | -5.41 | 0.718 | -7.54 | $4.81 \times 10^{-14}$ | *** |
| PGR | -31.2 | 3.85 | -8.11 | $4.89 \times 10^{-16}$ | *** |
| MD | -5.63 | 0.657 | -8.58 | $<2.00 \times 10^{-16}$ | *** |
| FG_Semelparous:PGR | 1.84 | 2.87 | 0.643 | 0.521 |  |
| FG_Shrub:PGR | 3.89 | 2.21 | 1.76 | 0.0788 | . |
| FG_Tree:PGR | -2.85 | 2.52 | -1.13 | 0.259 |  |
| FG_Semelparous:MD | -5.33 | 3.69 | -1.45 | 0.148 |  |
| FG_Shrub:MD | 3.22 | 1.90 | 1.70 | 0.0895 | . |
| FG_Tree:MD | 8.57 | 0.832 | 10.3 | $<2.00 \times 10^{-16}$ | *** |
| PGR:MD | 41.7 | 5.41 | 7.71 | $1.31 \times 10^{-14}$ | *** |

(2) Fecundity
|  | Estimate | Std. Error | z value | p value |  |
| --- | --- | --- | --- | --- | --- |
| (Intercept) | 1.70 | 0.537 | 3.16 | 0.00157 | ** |
| FG_Semelparous | -1.97 | 2.93 | -0.675 | 0.500 |  |
| FG_Shrub | -1.85 | 1.41 | -1.31 | 0.190 |  |
| FG_Tree | -3.33 | 0.696 | -4.78 | $1.73 \times 10^{-6}$ | *** |
| PGR | -14.6 | 4.58 | -3.20 | 0.00139 | ** |
| MD | -2.72 | 0.737 | -3.70 | $2.18 \times 10^{-4}$ | *** |
| FG_Semelparous:PGR | 2.50 | 2.71 | 0.921 | 0.357 |  |
| FG_Shrub:PGR | 2.16 | 2.17 | 0.995 | 0.320 |  |
| FG_Tree:PGR | 3.94 | 2.20 | 1.79 | 0.0732 | . |
| FG_Semelparous:MD | 3.10 | 4.37 | 0.709 | 0.479 |  |
| FG_Shrub:MD | 1.81 | 1.71 | 1.06 | 0.290 |  |
| FG_Tree:MD | 3.67 | 0.859 | 4.27 | $1.96 \times 10^{-5}$ | *** |
| PGR:MD | 27.6 | 6.08 | 4.55 | $5.47 \times 10^{-6}$ | *** |

(3) Growth
|  | Estimate | Std. Error | z value | p value |  |
| --- | --- | --- | --- | --- | --- |
| (Intercept) | 2.69 | 0.516 | 5.21 | $1.91 \times 10^{-7}$ | *** |
| FG_Semelparous | -2.88 | 3.38 | -0.851 | 0.395 |  |
| FG_Shrub | -2.79 | 1.55 | -1.79 | 0.0728 | . |
| FG_Tree | -5.28 | 0.703 | -7.50 | $6.33 \times 10^{-14}$ | *** |
| PGR | -21.3 | 4.80 | -4.44 | $8.87 \times 10^{-6}$ | *** |
| MD | -2.66 | 0.707 | -3.77 | $1.65 \times 10^{-4}$ | *** |
| FG_Semelparous:PGR | 4.75 | 2.67 | 1.78 | 0.0747 | . |
| FG_Shrub:PGR | 3.33 | 2.18 | 1.53 | 0.127 |  |
| FG_Tree:PGR | 7.63 | 2.15 | 3.55 | $3.82 \times 10^{-4}$ | *** |
| FG_Semelparous:MD | 4.02 | 5.01 | 0.802 | 0.423 |  |
| FG_Shrub:MD | 2.66 | 1.87 | 1.43 | 0.153 |  |
| FG_Tree:MD | 5.82 | 0.861 | 6.76 | $1.38 \times 10^{-11}$ | *** |
| PGR:MD | 34.0 | 6.25 | 5.44 | $5.45 \times 10^{-8}$ | *** |

\*\*\* $p < 0.001$ , \*\* $p < 0.01$ , \* $p < 0.05$ , . $p < 0.1$

## DISCUSSION

Patterns of interstage flow varied among plant functional types (Table 1, and Fig. 3) in ways that are consistent with life history tradeoffs and that are broadly similar to those based on elasticities of vital rates (Salguero-Gómez et al., 2015). However, interstage flow generally ascribed a larger role for fecundity than did elasticity, and that pattern was consistent across functional groups (Fig. 3). In addition, while growing populations tended to be dominated by interstage flows related to fecundity they tended to have high elasticities related to growth (Figs. 4, 5).

The different patterns we observed for interstage flow and elasticity reflect their different perspectives on population growth. Interstage flow is derived from the steady-state properties of populations (i.e., stable stage distributions and vital rates). Interstage flow, therefore, describes the current contribution of individual developmental stage transitions to population growth rate. Interstage flow is a retrospective analysis that describes how population growth rate has been influenced by events and conditions in the recent past. In contrast, elasticity is a prospective analysis that describes the relative contribution of vital rates to expected future population growth rate after a theoretical perturbation (Caswell, 2001).

Retrospective and prospective approaches provide complementary perspectives that are useful in different contexts (Horvitz et al. 1997). For example, elasticity analysis has been commonly used in the management of non-native invasive populations to predict how targeting control efforts at different life stages or vital rates will influence future population growth (Kerr et al. 2016). However, a retrospective approach is needed to understand how spatial variation in environmental conditions across the introduced range or periodic variation in conditions (such as associated with disturbance) affect the demography and growth of invasive populations. A common retrospective approach in these types of studies is life-table response experiments (LTRE), which describes the degree to which differences in population growth under different conditions reflect differences in the contribution made by each of the vital rates (Caswell 2010). For example, using field experiments Akin-Fajiye and Gurevitch (2020) found that disturbance significantly increased the population growth of invasive *Centaurea stoebe* (spotted knapweed). They then used LTRE to show that the increase in λ was overwhelmingly driven by increased fecundity. Interstage flow matrices could be a useful complement to LTRE analyses, by decomposing differences in λ among populations or treatments explicitly in terms of the contribution of individual flows between stages. Interstage flows represent stable stage–transition distributions, which are more useful than stable stage distributions because they can be decomposed in terms of transitions related to stasis, fecundity, and growth.

Such explicit descriptions of individual flows could be helpful in developing mechanistic stage structured demographic models that link the often density dependent dynamics of competitors or predators. Interstage flow may also be useful for addressing questions that involve the link between population dynamics and broader ecosystem properties. For example, plants allocate assimilated energy to processes that support fecundity, growth, and survival (Harper, 2010). The degree to which relative allocations vary across functional groups or across populations experiencing different environmental conditions can affect ecosystem processes. For example, when resource availability is low or environmental stressors such as herbivory and abiotic stress are high, most of the gross plant assimilated energy may be allocated to support functions related to survival (such as herbivore defense), making less primary production available to consumers (Fridley, 2017). Interstage flow is a more direct way to link demography and energy flow.

The interstage flow matrix, like other population statistics, is potentially influenced by factors related to statistical model design. For instance, interstage flows and elasticity are dependent on matrix dimension. In addition, categorizing matrix elements to different demographic processes involves a degree of ambiguity and discretion. For example, we classified interstage flow matrix elements relating to asexual reproduction (i.e., clonal growth) into growth, the same as transitions into larger size classes (i.e., growth of smaller plants into larger ones). However, these are potentially distinct demographic processes (Salguero-Gómez, 2018).

This proposed population statistic, the interstage flow matrix, describes the contribution of individual transitions between developmental stages to population growth. We suggest that future research use the interstage flow matrix to decompose population growth rate into population projection matrix elements in order to specify contributions to population growth rates.

## ACKNOWLEDGMENTS

This work was supported by the Japan Society for the Promotion of Science Grants-in-Aid for Scientific Research (KAKENHI) numbers: 20K06821, 19H03294, 26291087, and 15H04418. This research was also performed by the Environment Research and Technology Development Fund (JPMEERF20225005) of the Environmental Restoration and Conservation Agency provided by Ministry of the Environment of Japan. We are grateful to Hal Caswell, Hiroshi Tomimatsu, Masashi Kamo, and Fumie Tanaka for support and helpful comments.

## AUTHORS’ CONTRIBUTIONS

H.Y. and T.T. conceived the research question, developed the methodology, and wrote the original manuscript. H.Y. K.F. and T.T analyzed the data and conducted the statistical analysis. H.Y., K.F. and Y.K. developed the programming code for organizing the data. H.Y., K. F., J. L., and T.T. reviewed and edited the manuscript.

## DATA AVAILABILITY STATEMENT

The plant matrix database will be available via COMPADRE (https://compadre-db.org).

