## supporting information for "Interstage flow matrices: population statistic derived from matrix population models"

**Supplementary Material for "Interstage flow matrices: population statistic derived from matrix population models."**

[†] Deceased

APPENDIX A

*Sum of inter-stage flow matrix elements*

The sum of inter-stage flow matrix elements $f_{ij}^{\mathrm{IS}}$ was calculated as follows:

$$\sum_{i,j} f_{ij}^{\mathrm{IS}} = \mathbf{1}^{\mathrm{T}}\mathbf{A}\mathbf{w} = \mathbf{1}^{\mathrm{T}}\lambda\mathbf{w} = \lambda,$$

where $\mathbf{1}^{\mathrm{T}}\mathbf{w} = 1$ ($\mathbf{1} =$ vector of ones), and $\mathbf{A}\mathbf{w} = \lambda\mathbf{w}$. The sum of inter-stage flow matrix elements is therefore equivalent to the population growth rate.

### APPENDIX B

*Relationship between interstage flow and elasticity*

An interstage flow matrix element $f_{ij}^{\mathrm{IS}}$ is defined by $f_{ij}^{\mathrm{IS}} = a_{ij}w_j$, and the sum of the interstage flow matrix elements equals the population growth rate λ, i.e., $\lambda = \sum_{i,j} f_{ij}^{\mathrm{IS}}$ . (see Appendix A)

The definition of elasticity is as follows:

$$e_{km} = \frac{a_{km}}{\lambda}\frac{\partial\lambda}{\partial a_{km}} = \frac{a_{km}}{\lambda}\left(\sum_{i,j}\frac{\partial f_{ij}^{\mathrm{IS}}}{\partial a_{km}}\right), \qquad \text{(B1)}$$

$$\sum_{k,m} e_{km} = \sum_{k,m}\frac{a_{km}}{\lambda}\frac{\partial\lambda}{\partial a_{km}} = \sum_{k,m}\frac{a_{km}}{\lambda}\left(\sum_{i,j}\frac{\partial f_{ij}^{\mathrm{IS}}}{\partial a_{km}}\right) = 1, \qquad \text{(B2)}$$

where $\frac{\partial f_{ij}^{\mathrm{IS}}}{\partial a_{km}} = a_{ij}\frac{\partial w_j}{\partial a_{km}}$, if $i \neq k$ and $j \neq m$ and $\frac{\partial f_{ij}^{\mathrm{IS}}}{\partial a_{km}} = a_{ij}\frac{\partial w_j}{\partial a_{km}} + w_j$, if $i = k$ and $j = m$. Elasticity $e_{km}$ is the sum of the changes of the interstage flow $f_{ij}^{\mathrm{IS}}$ with respect to the change of population matrix elements multiplied by $a_{km}/\lambda$.
